# Rapid embryonic cell cycles defer the establishment of heterochromatin by Eggless/SetDB1 in *Drosophila*

**DOI:** 10.1101/450155

**Authors:** Charles A. Seller, Chun-Yi Cho, Patrick H. O’Farrell

## Abstract

Acquisition of chromatin modifications during embryogenesis distinguishes different regions of an initially naïve genome. In many organisms, repetitive DNA is packaged into constitutive heterochromatin that is marked by di/tri methylation of histone H3K9 and the associated protein HP1a. These modifications enforce the unique epigenetic properties of heterochromatin. However, in the early *Drosophila melanogaster* embryo the heterochromatin lacks these modifications which only appear later when rapid embryonic cell cycles slow down at the Mid-Blastula Transition or MBT. Here we focus on the initial steps restoring heterochromatic modifications in the embryo. We describe the JabbaTrap, a technique for inactivating maternally provided proteins in embryos. Using the JabbaTrap we reveal a major requirement for the methyltransferase Eggless/SetDB1 in the establishment of heterochromatin. In contrast, other methyltransferases contribute minimally. Live-imaging reveals that endogenous Eggless gradually accumulates on chromatin in interphase, but then dissociates in mitosis and its accumulation must restart in the next cell cycle. Cell cycle slowing as the embryo approaches the MBT permits increasing accumulation and action of Eggless at its targets. Experimental manipulation of interphase duration shows that cell cycle speed regulates Eggless. We propose that developmental slowing of the cell cycle times embryonic heterochromatin formation.

---

The transformation of the egg into a multicellular organism remains one of the wonders of the natural world. By the 18^th^ century, scientists viewed this process as an example of epigenesis, where the simple and formless fertilized egg progressively differentiates into a complex organized creature. We now know that the transformation of the egg from simple to differentiated is preceded and caused by a similar transformation of the naïve zygotic genome. The division of the genome into distinct euchromatin and heterochromatin is the most obvious example of genomic differentiation. Eukaryotes constitutively package significant portions of their genomes into heterochromatin (Brown 1966), and such packaging is critical for the stability of the genome (Janssen et al. 2018). Constitutive heterochromatin displays conserved genetic and molecular properties that are stably transmitted throughout development and across generations (Allshire and Madhani 2018). However, embryogenesis interrupts this stability, and the early embryos of many animals lack true heterochromatin. Thus, embryogenesis involves the restoration of heterochromatin, and this restoration can serve as a paradigm for how epigenetic control of the genome arises during development.

In general, constitutive heterochromatin is transcriptionally silent, late replicating, and low in recombination. The chromosomes of many species contain large megabase sized arrays of simple repeated sequences, known as satellite DNA, surrounding their centromeres, and it is this pericentric repetitive DNA that composes the bulk of the constitutive heterochromatin. Although heterochromatin was originally defined cytologically – it remains condensed throughout the cell cycle – modern studies often emphasize the set of conserved modifications to its chromatin. Heterochromatic nucleosomes are hypoacetylated and di/tri-methylated at lysine 9 on the N-terminal tail of Histone H3 (hereafter H3K9me2/3). The deposition of H3K9me2/3 creates a binding site for HP1 proteins (Eissenberg and Elgin 2014) and HP1 can oligomerize and recruit additional heterochromatin factors which could then generate a distinct compartment in the nucleus.

Studies in *Drosophila* have historically made major contributions to our understanding of heterochromatin (Elgin and Reuter 2013). Many key regulators of heterochromatin were discovered in genetic screens for modifiers of Position Effect Variegation (PEV), a phenomenon where a reporter gene close to the border between euchromatin and heterochromatin is differentially silenced among different cells in an all or none fashion. Currently over 150 distinct genetic loci are reported to impact PEV, highlighting the complexity of heterochromatin control. Critically, the pathway depositing H3K9me2/3-HP1a forms the regulatory core and null mutations in *Su(var)2-5* (HP1a), and replacement of H3K9 with arginine (H3K9R) to block its modification are both lethal (Eissenberg et al. 1990; Penke et al. 2016, 2018). The *Drosophila* genome encodes three K9 methyltransferases: *Su(var)3-9, eggless/SetDB1, and G9a*, and all three have been reported to modify PEV (Mis et al. 2006; Brower-Toland et al. 2009). To our knowledge, no study has systematically assessed the contributions of these enzymes to the establishment of heterochromatin in the embryo.

Although a dispensable gene (Tschiersch et al. 1994), *Su(var)3-9* is the best studied and most strongly associated with the constitutive heterochromatin in the literature (Elgin and Reuter 2013; Allshire and Madhani 2018). Loss of *Su(var)3-9* leads to strong suppression of PEV. Additionally, analysis of salivary gland polytene chromosomes revealed that *Su(var)3-9* is required for the localization of H3K9me2/3 and HP1a to the chromocenter, the region where the pericentric heterochromatin clusters in the nucleus (Schotta et al. 2002). G9a is dispensable in *Drosophila* and thought to function primarily at euchromatic sites (Seum et al. 2007b). Eggless is the only essential K9 methyltransferase. Studies in larval and adult stages have emphasized that egg plays a specialized role in depositing H3K9me2/3 at Chromosome 4 (Seum et al. 2007a; Tzeng et al. 2007) which forms a distinct chromatin structure (Riddle et al. 2012). H3K9me2/3-HP1a is missing from chromosome 4 in the polytene chromosomes of egg mutants, but unaffected at the chromocenter. These findings suggested that egg does not regulate the pericentric heterochromatin. Further analysis demonstrated that egg is also required in both the soma and germline for the completion of oogenesis (Clough et al. 2007, 2014; Wang et al. 2011). Egg plays a major role in the piRNA pathway that silences transposons in the germline (Rangan et al. 2011; Sienski et al. 2015; Yu et al. 2015). Although they have yielded much information, these studies focused on the maintenance of heterochromatin, but not on its embryonic emergence.

Like most animals, the *Drosophila* embryo begins development with a stereotyped series of rapid cell cycles following fertilization that are driven by high levels of cyclin-CDK1 activity (Farrell and O’Farrell 2014; Yuan et al. 2016). These divisions occur in a syncytium, are synchronous, lack gap phases, and run entirely by maternal gene products. The first 8 divisions are remarkably fast (each with an interphase of less than 4 min) and occur deep within the embryo. Starting at cycle 9, nuclei migrate to the surface of the embryo to form a syncytial blastoderm. Interphase duration extends progressively and incrementally from cycle 10 through 13 (from 7 min to 14 min). Beginning at cycle 14 the embryo enters the Mid-Blastula Transition, or MBT, a conserved embryonic transformation. At the MBT, the cell cycle slows down dramatically due to the programmed downregulation of CDK1, and the embryo switches its focus from cell proliferation to morphogenesis and patterning. Concomitantly, the zygotic genome activates in full and stable patterns of histone modification emerge (Chen et al. 2013; Li et al. 2014). Zygotic transcription then drives morphogenetic events like cellularization and gastrulation. The slowing of the cell cycle is critical for the developmental events that follow it because short cell cycles inhibit productive transcription (Shermoen and O’Farrell 1991; Strong et al. 2017). Additionally, rapid cell cycles could impair the installation of chromatin modifications that regulate gene expression, but to our knowledge no direct examination of the enzymes installing these modifications at the MBT exists.

Prior work from our lab has outlined a series of changes made to the heterochromatin as the embryo reaches the MBT. Surprisingly, the conserved features of heterochromatin appear at different times during embryogenesis, indicating that diverse mechanisms contribute to the introduction of its otherwise well correlated properties. The compaction of satellite DNA is evident as early as cycle 8, before the onset of late replication and the association of H3K9me2/3-HP1a (Shermoen et al. 2010). Upon Cdk1 downregulation at the MBT select satellite repeats then recruit the protein Rif1 which acts to delay their replication (Seller and O’Farrell 2018). During the prolonged interphase 14 some satellites become stably associated with H3K9me2/3-HP1a (Yuan and O’Farrell 2016), and by the cycle 15 the satellites coalesce into a clear chromocenter. The emergence of the key features of heterochromatin follows a stereotyped schedule centered around the MBT, and we suggest that this period represents the establishment of mature heterochromatin in the embryo. Following the MBT the satellites faithfully exhibit these characteristics for the rest of development. We want to understand the developmental control of establishment, but first need to discover the key molecules in this process. Here we identify the methyltransferase Egg as a key initiator and propose that heterochromatin formation is timed by the limitations imposed on Egg by rapid embryonic cell cycles.

## RESULTS

### The JabbaTrap inactivates maternally deposited proteins by rapid mislocalization

The power of fly genetics is nearly incontestable, but the early embryo does present a significant challenge to it. Over 80% of genes are supplied to the embryo as maternally deposited mRNA and protein, and consequently development of the embryo up to the MBT does not require zygotic gene products. Removing maternal contributions is of course possible when the mother fly is mutant for the gene under study, but many interesting genes have essential zygotic functions. Germline clone techniques can serve as a workaround for the study of essential genes, but only if the mutation under study does not interfere with oogenesis. Here the genetics of egg serves as a clear example of the dilemma. Zygotic null mutants for egg have poor viability, and egg is required at multiple points in both the soma and germline during oogenesis. Therefore, no study has examined a potential role for egg during the establishment of the heterochromatin in the early embryo.

Alternative strategies for removing maternally supplied function are needed. Although some methods do exist, they depend on the stability of their targets and can be slow and incomplete. Here we developed an approach in the spirit of the Anchors Away technique (Haruki et al. 2008) for inactivating GFP tagged proteins in the early embryo by selective mislocalization. As schematized in Fig 1A, target proteins are mislocalized by the expression of an anti-GFP nanobody (Rothbauer et al. 2008) fused to the lipid droplet binding protein Jabba (Li et al. 2012), which we call the JabbaTrap. Lipid droplets act as storage organelles during early embryogenesis and are distributed throughout the cytoplasm (Welte 2015), and we reasoned that trapping nuclear proteins on lipid droplets would block their function.

**Fig. 1.**
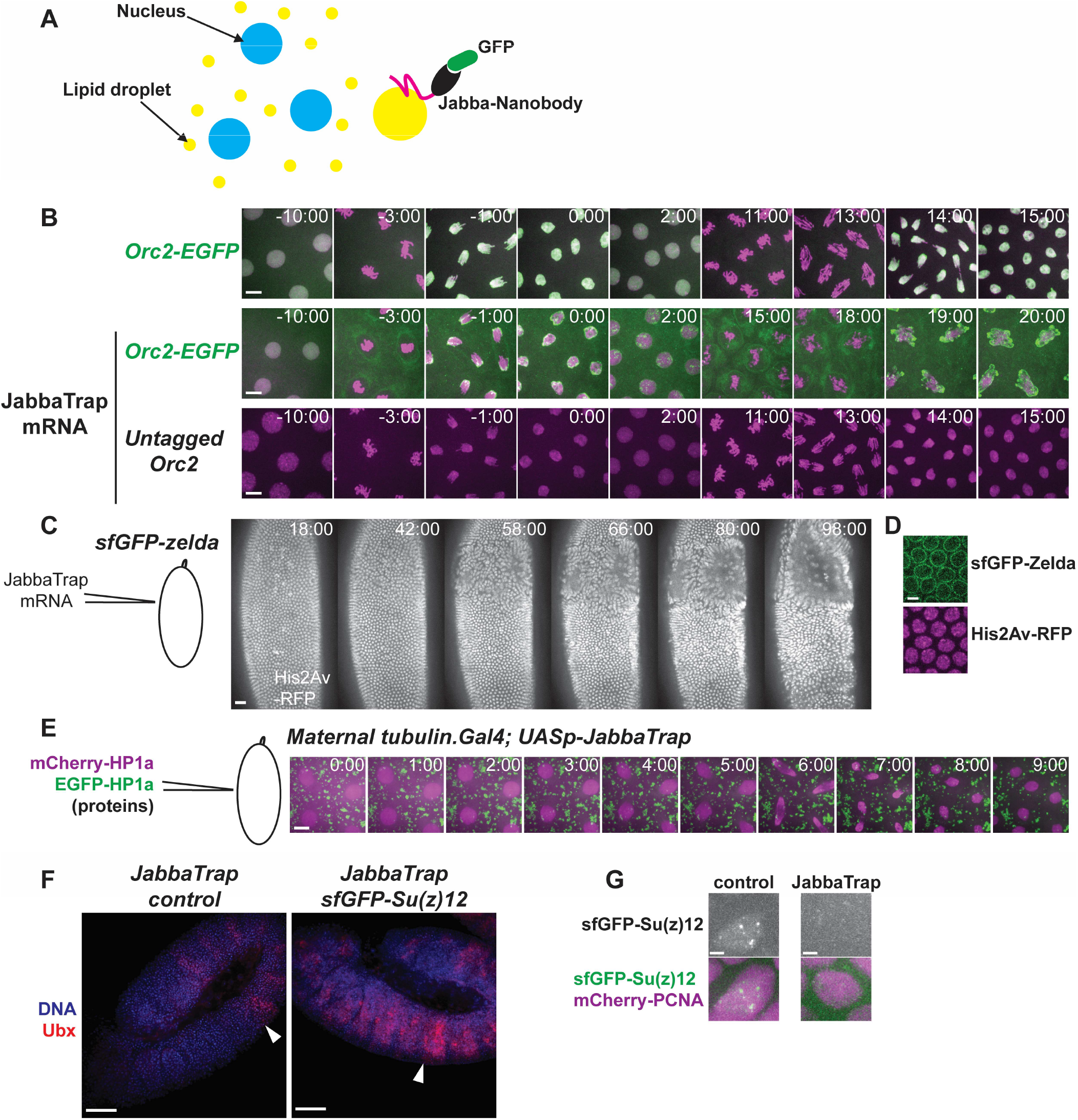
JabbaTrap rapidly mis-localizes and blocks nuclear proteins in the embryo. **A)** Schematic of the JabbaTrap technique. Lipid droplets (yellow circles), which serve as storage organelles, are distributed throughout the cytoplasm of the syncytial embryo. Ectopic expression of Jabba fused to an anti-GFP nanobody traps GFP tagged proteins on lipid droplets. **B)** The Jabbatrap mislocalizes Orc2 and blocks DNA replication. Stills from live imaging of embryos of the indicated genotype after injection of Cy5 labeled histone proteins alone or together with JabbaTrap mRNA. Top row, Orc2-GFP embryos injected with labeled histones (shown in magenta) only. Orc2 (shown in green) rapidly binds to the anaphase chromosomes (−1:00) and remains nuclear during the subsequent interphase. In Orc2-GFP embryos injected with both labeled histones and Jabbatrap message (middle row), Orc2 is prevented from binding to the chromosomes and mislocalized to both the reforming nuclear envelope and to puncta in the cytoplasm. Anaphase bridging occurs in the following mitosis (19:00). Bottom row, wildtype embryos injected with both Cy5-histones and JabbaTrap mRNA undergo normal mitotic cycles. Scale bar is 3 µm. See Supplemental Movie 1. **C)** Stills from live imaging following His2Av-RFP in an sfGFP-Zelda embryo injected with JabbaTrap mRNA. Nuclei near the injection site failed to cellularize and collapsed out of the cortical layer. Scale bar is 10 µm and the start of cycle 14 is set as 0:00. **D)** Single z plane showing the mislocalization of sfGFP-Zelda by expression of JabbaTrap mRNA. Scale bar is 2 µm. **E)** Co-injection of mCherry-HP1a and GFP-HP1a into an embryo from mothers expressing transgenic JabbaTrap during oogenesis. Scale bar is 4 µm. **F)** JabbaTrap expression blocks sfGFP-Su(z)12 leading to derepression of the homeotic protein Ubx. The white arrowhead indicates the wild-type anterior border of Ubx expression. Embryos stained with DAPI (blue) and anti-Ubx (red). Scale bar is 25 µm. **(G)** Control or JabbaTrap expressing *sfGFP-Su(z)12* embryos injected with mCherry-PCNA to label the nucleus. The JabbaTrap mislocalizes Su(z)12 to the cytoplasm. Scale bar is 1 µm.

As an initial test, we targeted the protein Orc2, an essential component of the Origin Recognition Complex (ORC). Orc2 has a highly reproducible pattern of localization during the cell cycle and is required for S phase completion in mitotic cycles (Baldinger and Gossen 2009). Orc2-GFP embryos were injected with either water (control) or synthesized JabbaTrap mRNA and then immediately filmed by confocal microscopy. In control injected embryos Orc2-GFP resided in the nucleus in interphase, diffused into the cytoplasm upon mitotic entry, and then rapidly bound along the anaphase chromosomes in preparation for the next S phase (Fig. 1B, first row). In contrast, in embryos expressing the JabbaTrap, Orc2-GFP displayed aberrant localization. During the first mitosis following injection, Orc2 was present on numerous cytoplasmic puncta as well as on membranous structures surrounding the mitotic spindle (Fig. 1B, second row, 3:00, Supplemental Movie 1). Upon mitotic exit Orc2 coalesced around the reforming nuclear envelope but did not bind to the chromosomes and remained in the cytoplasm during the subsequent S phase. As expected, without chromatin bound Orc2 nuclei entered mitosis after a delay but then underwent catastrophic anaphase bridging (Fig. 1B, second row, 19:00). Importantly, expression of the JabbaTrap in embryos with untagged Orc2 did not disrupt cell cycle progression (Fig. 1B, third row).

As a second test we used the same approach to mislocalize the protein Zelda, a pioneering transcription factor required for proper zygotic genome activation (Liang et al. 2008). We injected *sfGFP-zelda; His2Av-RFP* embryos with JabbaTrap mRNA during nuclear cycle 10, and then followed them by time lapse microscopy. Zelda is a nuclear protein that forms foci on interphase chromatin. Expression of the JabbaTrap blocked the nuclear localization of Zelda near the injection site (Fig. 1D). Embryos with JabbaTrapped Zelda failed dramatically at cellularization and gastrulation, events known to require zygotic transcription. Nuclei near the injection site did not undergo the shape changes characteristic of cellularization, but rather appeared swollen and ultimately fell out of the cortical layer (Fig. 1C).

Next, we established transgenic lines for the JabbaTrap. Lipid droplets form during oogenesis and we reasoned that by driving the expression of the JabbaTrap with a Maternal-tubulin Gal4, which activates transcription after egg chamber budding, embryos would already have the JabbaTrap deployed when they begin developing. To test this idea, we injected purified mCherry-HP1a and GFP-HP1a proteins into embryos collected from mothers expressing the JabbaTrap. Following protein injection, we immediately filmed embryos by confocal microscopy. As expected mCherry-HP1a quickly localized to the nucleus after its delivery (Fig. 1E, in magenta). In contrast, injected GFP-HP1a localized to numerous cytoplasmic puncta and was excluded from the nucleus throughout the cell cycle (Fig. 1E, in green).

As a further test we used our transgenic JabbaTrap to block the Polycomb group protein *Su(z)12*, an essential component of PRC2 with well characterized mutant phenotypes at different stages of development. Polycomb group proteins are required for the repression of the homeotic genes and determine their anterior boundary of expression in the developing embryo. However, zygotic null mutants for *Su(z)12* die during L1 or L2 stages without any obvious polycomb phenotypes, likely due to the confounding impact of maternal supply (Birve et al. 2001). Germline clone analysis using null alleles revealed that *Su(z)12* is also essential for oogenesis. To test an embryonic function for *Su(z)12*, Birve et al. crossed females with germline clones of the weak *Su(z)12^2^* allele with males heterozygous for a deficiency spanning the gene region. The resulting embryos misexpressed the homeotic gene *Ubx*, a classic polycomb phenotype. To determine if our JabbaTrap approach could produce a similar loss of function situation we tagged Su(z)12 with sfGFP at its endogenous locus using CRISPR-Cas9 (Supplemental Fig. 1) and expressed our JabbaTrap maternally in this background. This resulted in high embryonic lethality (2.7% hatch rate, n = 150) and clear misexpression of *Ubx* (Fig. 1F). In control embryos Su(z)12 formed nuclear foci in gastrulating embryos – a stage when polycomb bodies first become evident. In contrast, expression of the JabbaTrap prevented nuclear localization of Su(z)12 (Fig. 1G)

### Eggless is required for the establishment of the heterochromatin at the MBT

Our prior study characterized the appearance of H3K9me2/3 during early embryogenesis using antibody staining and observed little evidence of this modification before cycle 13 (Yuan and O’Farrell 2016). To understand the dynamics of H3K9me2 deposition we took advantage of the Fab-based live endogenous modification (FabLEM) technique (Hayashi-Takanaka et al. 2011). We injected GFP-HP1a expressing embryos with Cy5 labeled Fab fragments recognizing H3K9me2 to follow both modifications in real time over cycles 12-14 (Fig. 2A, Supplemental Movie 2). In doing so we made several interesting observations. Within a cell cycle, the K9me2 Fab accumulated progressively in chromatin foci as interphase proceeded, but the signal dissipated upon entry into mitosis. At telophase the K9me2 Fab was again recruited to chromatin, and in each successive cycle the intensity of this telophase signal increased. We interpret this as an indication of the amount of H3K9me2 carried over from the previous cell cycle, and by cycle 14 this telophase signal is significant (Fig. 2A, Cycle 14 0:00). Additionally, as interphase duration increased in successive cycles the chromatin bound Fab signal accumulated to greater extent. This was particularly dramatic during interphase 14 when the K9me2 Fab signal became quite bright and eventually covered the chromocenter (Fig. 2A, Cycle 14 64:00). The recruitment of HP1a to chromatin foci closely paralleled the appearance of K9me2 Fab signal. We conclude that H3K9me2 increases progressively during cycles 12-14 as interphase duration extends, and that modifications deposited in earlier cycles can carry over to later cycles.

**Fig. 2.**
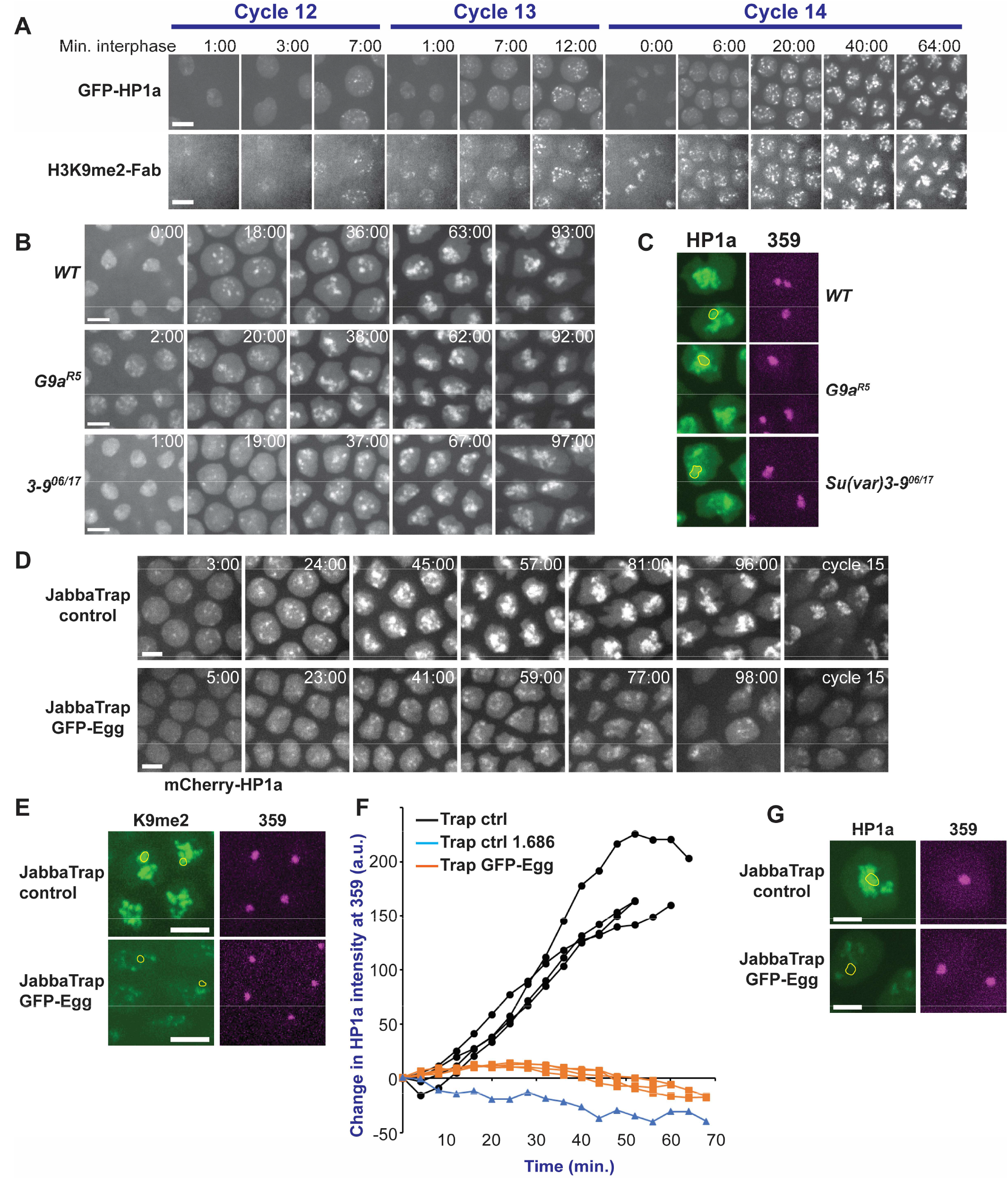
Eggless is required for the embryonic establishment of heterochromatin. **A)** Progressive accumulation of H3K9me2-HP1a in the early embryo. Selected stills from live imaging over 3 cell cycles of GFP-HP1a embryos injected with Cy5 labeled H3K9me2 Fab fragment. Scale bar is 3 µm. See Supplemental Movie 2. (B,**C)** G9a and Su(var)3-9 are not required for the establishment of heterochromatin. **B)** Selected stills from live imaging of embryos of the indicated genotype injected with mCherry-HP1a protein. The beginning of interphase 14 was set as 0:00. The scale bar is 2 µm. **C)** Stills of live embryos of the indicated genotype injected with GFP-HP1a and 359-TALE-mCherry. The location of 359 is outlined in yellow on the HP1a channel. Images were acquired approximately 1 hour into interphase 14. **(D-G)** JabbaTrapping Egg blocks the embryonic onset of H3K9me2-HP1. **D)** Live imaging of mCherry-HP1a injected into wildtype control or GFP-Egg embryos following maternal expression of the JabbaTrap. Mislocalization of Egg significantly reduced and delayed the accumulation of HP1a at the heterochromatin. The start of interphase 14 was set as 0:00. The scale bar is 3 µm. Compare Supplemental Movies 3 and 4. **E)** Stills from live embryos of the indicated genotype injected with Cy5 labeled H3K9me2 Fab fragments and 359-TALE-mCherry. The position of 359 is outlined in yellow on the H3K9me2 channel. Images were acquired approximately 1 hour into interphase 14. Scale bar is 2 µm. **F)** Plot of the change in mCherry-HP1a fluorescence at the 359 bp repeat over time during interphase 14. The 1.686 satellite, which does not recruit HP1a, serves as a negative control. Each line represents a different embryo, where fluorescence was measured in over 20 nuclei per embryo and the mean average for each time point was calculated. **G)** Stills from live embryos of the indicated genotype injected with mCherry-HP1a and 359-TALE-HaloJF646. The position of 359 is outlined in yellow on the HP1a channel. Embryos are approximately 80 minutes into interphase 14, during cephalic furrow formation. The scale bar is 1 µm.

Because the association of HP1a during interphase 14 tracked closely the accumulation of H3K9me2 we used its recruitment to assess the contributions of the different *Drosophila* K9 methyltransferases to the establishment of heterochromatin. As mentioned above, *G9a* and *Su(var)3-9* are dispensable genes, allowing us to analyze embryos from null backgrounds. Embryos of either *WT, G9a^RG5^*, or *Su(var)3-9^06/17^* background were injected with mCherry-HP1a and the recruitment of this protein to heterochromatin was recorded during cycle 14 by confocal microscopy. The timing and extent of HP1a recruitment to heterochromatic foci appeared similar between *WT* and *G9a^RG5^* embryos, and by the end of cycle 14 these foci had fused into a chromocenter (Fig. 2B, 2^nd^ row). Surprisingly, although HP1a recruitment in *Su(var)3-9* embryos was altered, with both delayed and reduced appearance of foci, these embryos still formed significant amounts of heterochromatin (Fig. 2B, 3^rd^ row). TALE-light probes allow us to follow specific satellite repeats in live embryos (Yuan et al. 2014). Using this technique, we previously documented that the 359 bp repeat, a 20 Mb sized satellite on the X chromosome, acquires H3K9me2/3-HP1a during cycle 14. By co-injecting mCherry-HP1a and 359-TALE-GFP proteins we observed that neither *G9a* or *Su(var)3-9* was required for the recruitment of HP1a at 359 (Fig. 2C). We conclude that *G9a* is not required for the establishment of heterochromatic modifications, and that *Su(var)3-9*, while contributing, is not required for initiating HP1a recruitment.

Our development of the JabbaTrapping technique allowed us to assess the contributions of eggless to establishment. First, we used CRISPR-Cas9 to tag endogenous Egg with GFP at its N-terminus (Supplemental Fig. 1), the resulting line was healthy and fertile indicating that our tag is functional. In JabbaTrap expressing embryos GFP-Egg was mislocalized to small cytoplasmic puncta and blocked from entering the nucleus throughout the cell cycle (Supplemental Fig. 2). Like the behavior of lipid droplets, these puncta were prominent and surrounded the nuclei during the syncytial cycles but appeared to drop out of the cortical layer by cycle 14. Additionally, embryos with JabbaTrapped Egg had poor viability (15% hatch rate, n = 283).

To assess the effect that mislocalization of Egg had on the establishment of the heterochromatin JabbaTrap control and JabbaTrapped GFP-Egg embryos were injected with mCherry-HP1a and imaged by confocal microscopy during cycle 14. In control embryos HP1a formed foci that grew in number and intensity over time, and then ultimately fused into a large chromocenter. Once established during cycle 14, HP1a association at the chromocenter is inherited in subsequent cell cycles, and nuclei begin interphase 15 with HP1a enriched at the chromocenter (Fig. 2D, Supplemental Movie 3). In contrast, the mislocalization of Egg substantially reduced and delayed the recruitment of HP1a to heterochromatic foci (Fig. 2D, Supplemental Movie 4). Even by the end of cycle 14 the amount of HP1a marked heterochromatin was clearly less than control, indicating that some of the sequences that normally acquire HP1a fail to do so in the absence of Egg. Indeed, following JabbaTrapping of Egg, HP1a did not localize to the 359 bp repeat (Fig. 2G). In control embryos, HP1a was recruited to the 359 bp repeat during the middle of interphase 14, but JabbaTrapping of Egg blocked this accumulation (Fig. 2F and Supplemental Fig.3). Additionally, the deficit in HP1a localization to the chromocenter was inherited into cycle 15 (Fig. 2D).

**Fig. 3.**
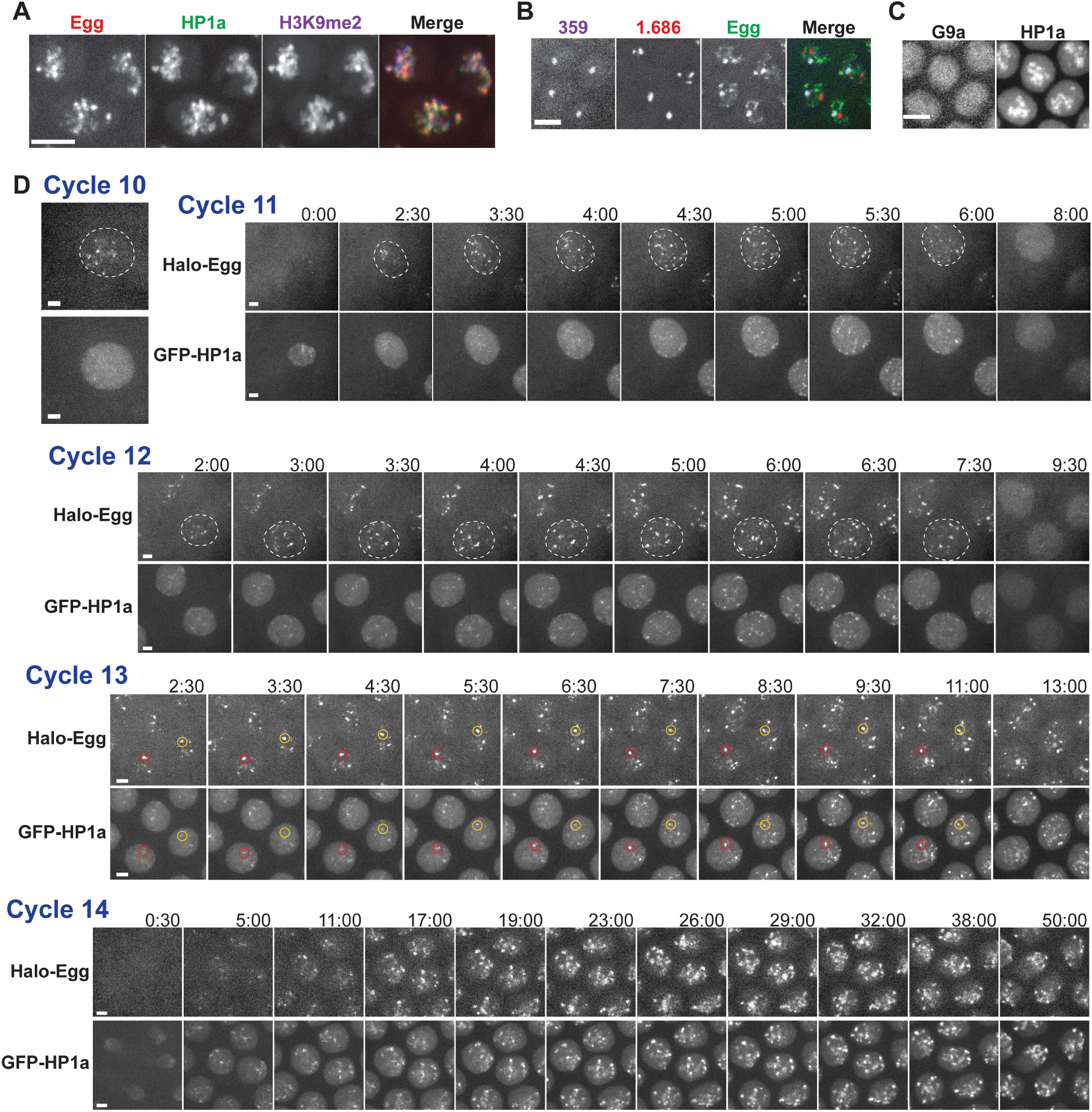
localization and dynamics of Egg during the establishment of heterochromatin. **A)** Confocal micrographs of a cycle 14 Halo(JF549)-Egg embryo injected with GFP-HP1a and Cy5 labeled H3K9me2 Fab. Scale bar 2 µm. **B)** Confocal micrograph of a cycle 14 GFP-Egg embryo injected with Halo(JF646)-359 and mCherry-1.686 TALE-Lights. Scale bar is 2 µm. **C)** Confocal micrographs of a live sfGFP-G9a embryo injected with mCherry-HP1a show that G9a does not localize to heterochromatin. Images were acquired approximately 50 min. into interphase 14. Scale bar 2 µm. **D)** Selected stills from live imaging of Halo(JF549)-Egg and GFP-HP1a during cycles 10-14 as interphase duration naturally extends. The recorded cycle is indicated above the micrographs. Egg accumulates at multiple bright foci in the interphase nucleus (denoted in dashed white line). Note that the recruitment of Egg precedes that of HP1a during development and during the cell cycle. Longer interphases permit increased accumulation of Egg on chromatin. Yellow and red circles in cycle 13 records track the fate of two foci of Egg and the corresponding locations in the HP1a channel. Each cell cycle was recorded from a different Halo-Egg embryo labeled with JF549 and the beginning of interphase was set as 0:00. Scale bars are 1 µm. See Supplemental Movie 5.

Next, we directly examined K9 methylation using the K9me2-Fab. JabbaTrapping Egg completely prevented the appearance of H3K9me2 at 359 during cycle 14, and significantly reduced its accumulation throughout the chromocenter (Fig. 2E). However, we note that the mislocalization of Egg did not completely abolish the formation of either HP1a or H3K9me2-Fab foci, these could represent sequences that rely on methylation by G9a or Su(var)3-9, or that recruit HP1a by a methylation independent pathway. To confirm these results, we stained fixed gastrulation stage embryos for HP1a and H3K9me3. Control embryos had a clear chromocenter that stained positively for both modifications. In contrast, in JabbaTrapped Egg embryos K9me3 was undetectable, and although a few HP1a foci were present, they were faint and disperse (Supplemental Fig. 3C).

We conclude that Egg plays a major role in the restoration of repressive chromatin modifications on the satellite DNA during the MBT and is required for the establishment of H3K9me2/3-HP1a at the 359 bp repeat. In addition, Egg is unique in this respect among the three K9 methyltransferases in *Drosophila*.

### Live imaging of Egg reveals the dynamics of heterochromatin establishment

Although prior studies had documented enrichment of Egg at Chromosome 4 and not to the chromocenter, our JabbaTrap experiments revealed a major role for Egg in the formation of the constitutive heterochromatin. We thus decided to examine the localization dynamics of Egg by live confocal microscopy in the early embryo. However, endogenous levels of Egg were relatively low in the embryo making long term and high-resolution imaging of GFP-Egg difficult. To solve this issue, we took advantage of the HaloTag system in combination with the JF549 fluorescent dye which has improved brightness and photostability compared to EGFP (Grimm et al. 2015). We used CRISPR-Cas9 to insert the HaloTag at the N-terminus of the endogenous egg gene. The resulting flies were viable and fertile. After permeabilization with Citrasolv, the HaloTag ligand JF549 could directly enter the embryo allowing easy *in vivo* labeling. Examination of embryos with both Halo549-Egg and GFP-Egg revealed that the two proteins had equivalent localization during late cycle 14 (Supplemental Fig. 4D). Additionally, the JF549 labeling was specific to embryos containing the Halotag (Supplemental Fig. 4E).

**Fig. 4.**
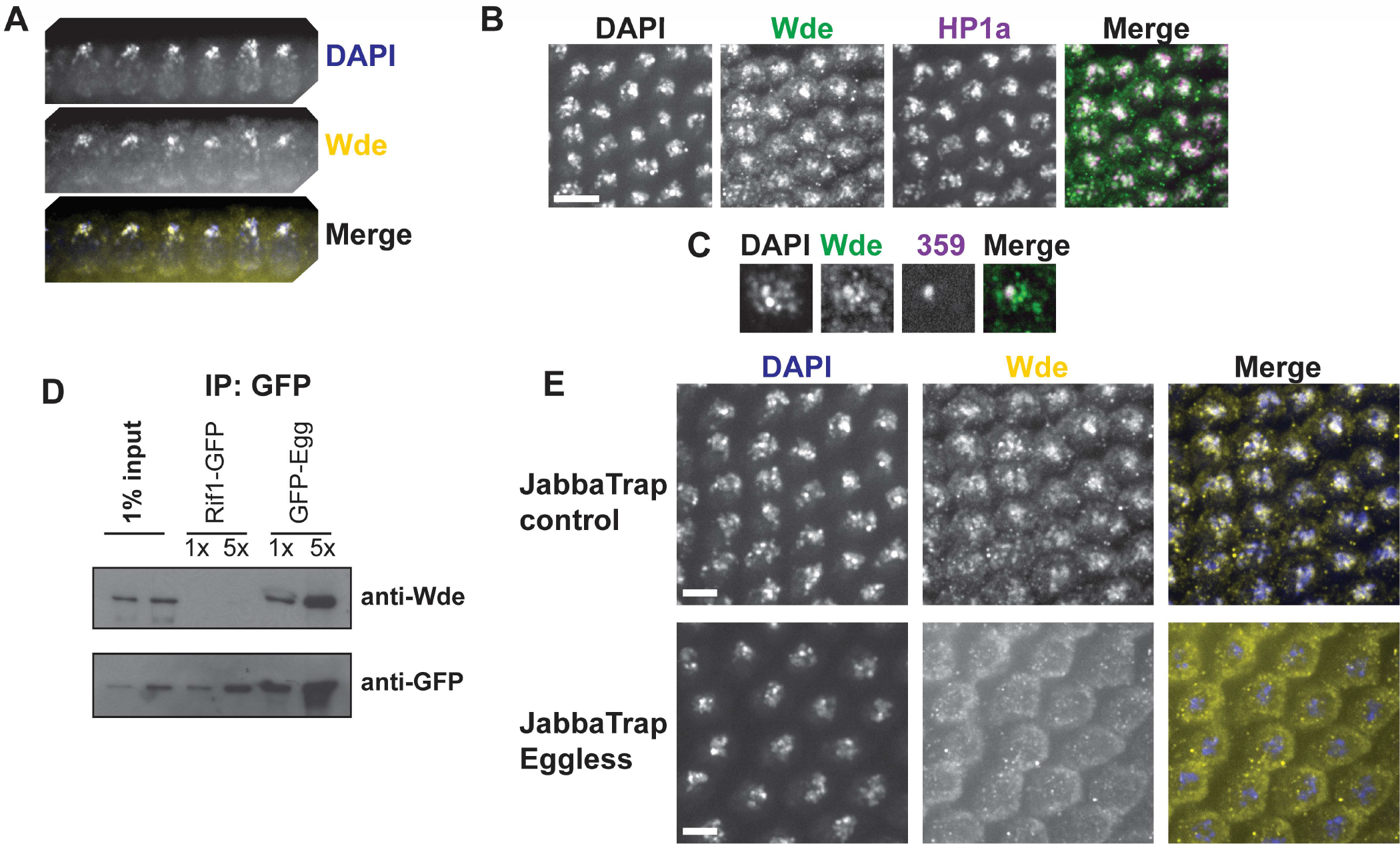
Windei, a cofactor of Egg, localizes to the heterochromatin in the embryo and interacts with Egg. **(A-C)** Wde is enriched at the constitutive heterochromatin in early embryos. **A)** Co-localization of Wde with the DAPI intense apical portion of the nucleus in fixed embryos. **B)** Cycle 14 embryos immunostained for HP1a and Wde. Scale bar is 5 µm. **C)** Fixed cycle 14 embryo stained for Wde and with a TALE-light recognizing the 359 bp repeat. **D)** GFP-Egg was immunoprecipitated from 1 to 3 hour old embryos and analyzed by western blotting. Wde interacts with Egg in the early embryo. **E)** Fixed embyros expressing the JabbaTrap in either a control or GFP-Egg background were immunostained for Wde. Wde is mislocalized to the cytoplasm when GFP-Egg is JabbaTrapped. Scale bar is 3 µm.

In late cycle 14 embryos Halo-Egg colocalized with GFP-HP1a and the H3K9me2-Fab at the satellite sequences (Fig. 3A). Interestingly, the Egg signal was contained within a broader nuclear region labeled with HP1a and K9me2 indicating that at this stage, when the heterochromatin has matured, Egg does not fill the entire heterochromatin domain. By altering the parental origin of tagged eggless we determined that the Egg present during cycle 14 is primarily maternally supplied (Supplemental Fig. 4C). Not all satellite sequences acquire H3K9me2/3 during the MBT, the 1.686 repeat remains free of this modification until much later in development (Yuan and O’Farrell 2016). In accordance with this fact, by using different TALE-light probes we see that Egg binds to 359, but not to 1.686 (Fig. 3B). Eggless contains a tandem tudor domain (Supplemental Fig. 4A), a structure known to mediate chromatin targeting in other proteins. Therefore, we generated a transgene to express a version of GFP-Egg lacking the tandem tudor sequence and expressed this mutant protein in the early embryo. We observed that the tandem tudor domain was required for the nuclear localization of Egg (Supplemental Fig. 4B). For comparison we also examined the localization of the G9a methyltransferase because our previous experiments demonstrated that it was not required for heterochromatin establishment. Unlike Egg, G9a did not co-localize with HP1a during cycle 14, but rather was diffusely present in the nucleus (Fig. 3C).

As early as cycle 10, we documented multiple bright foci of Halo-Egg in the nucleus (Fig. 3D). In contrast, at this stage HP1a is uniformly distributed in the nucleus and not enriched to heterochromatic foci. Live imaging of Halo549-Egg together with GFP-HP1a over the next 4 cell cycles yielded several interesting observations (Fig. 3D). During each interphase Egg accumulated progressively to bright chromatin foci after a short (2 min.) delay following exit from the prior mitosis (Fig. 3D, Supplemental Movies 5 and 6). By tracking individual Egg foci over multiple frames of our records we documented a pattern (Fig. 3D, circled foci, and Supplemental Movie 5) where a small focus of Egg would appear and then be joined by a focus of HP1a within minutes. The two proteins then grew in intensity over the course of interphase. Upon entry into mitosis, HP1a dissociated from chromatin and Egg followed slightly later. Overall, the recruitment of Egg to specific foci preceded that of HP1a within the cell cycle and during development. The longer interphase durations in later cell cycles correlated with greater accumulation and longer retention of Egg at its chromatin targets. For instance, during the brief 7 min. interphase 11 Egg foci grew in number and intensity for 5.5 min., but then these foci rapidly disappeared as the nucleus prepared for mitosis (Fig. 3D). In contrast, during the prolonged interphase 14 Egg foci were brighter and remained on chromatin for the duration of our records. In conclusion, the recruitment of Egg to its chromatin substrates is a progressive process occurring in each cell cycle that begins early in embryogenesis (at least by cycle 10). These observations together with our real time records of H3K9me2 accumulation (Fig. 2A) indicate that changes in interphase duration may time the formation of the heterochromatin.

Our records also revealed some diversity in behaviors. The appearance and growth of Egg foci did not follow a single schedule, apparently different sequences recruit the protein at different times during interphase. By cycle 14, it was clear that HP1a was recruited to some foci before Egg, a possible indication of the H3K9me2/3 carried over from prior cell cycles. Additionally, some foci of HP1a did not co-localize with Egg, which is consistent with our observation that a small amount of HP1a was still recruited after JabbaTrapping Egg (Fig. 2D).

### Windei, an activator of Eggless, localizes to the heterochromatin in the early embryo

Studies on both Egg, and its mammalian counterpart SETDB1 suggest that it requires a cofactor for efficient methylation of H3K9. SETDB1 alone has very poor methyltransferase activity (Schultz et al. 2002), but it is greatly stimulated when in complex with the protein mAM/MCAF1 (Wang et al. 2003). The Drosophila ortholog of mAM known as *windei (wde)* was suggested to play a similar role in flies (Koch et al. 2009). The phenotypes of *wde* and *egg* mutants are similar, and *wde* is also required during oogenesis. The two proteins physically interact and were reported to co-localize at Chromosome 4 in polytene preparations, but both were absent from the chromocenter. Because we have documented a role for Egg at the constitutive heterochromatin in the early embryo we decided to examine Wde during this stage.

Immunostaining revealed that Wde localizes to the heterochromatin during its establishment at the MBT. Wde was enriched in the DAPI intense apical portion of the nucleus where the satellite sequences reside (Fig. 4A). Additionally, we observed broad colocalization between Wde and HP1a late in cycle 14 (Fig. 4B). Because we had shown that the association of H3K9me2/3-HP1a to the 359 bp repeat required Egg we wanted to know if Wde also bound to this satellite. TALE-light staining revealed that Wde does localize to 359 (Fig. 4C). By co-IP western blotting we documented that GFP-Egg interacts with endogenous Wde in the early embryo (Fig. 4D), in agreement with what was reported in S2 cells (Koch et al. 2009). We also noticed that Wde was found in the cytoplasm in embryos with JabbaTrapped Egg (Fig. 4E), indicating that the JabbaTrap may be useful to mislocalize entire protein complexes. We conclude that in the early embryo Wde co-localizes with Egg during the formation of the constitutive heterochromatin.

### Interphase duration controls Egg recruitment to and establishment of the heterochromatin

To examine the relationship between heterochromatin establishment and the cell cycle more directly we injected Halo-Egg embryos with Cy5 labeled histone proteins, to visualize the chromatin, and imaged at high frame rate. As described previously, Egg accumulated to multiple nuclear foci during interphase with a slight delay following exit from the previous mitosis (Supplemental Movie 6). These foci abruptly disappeared upon chromosome condensation at prophase (Fig. 5A, Supplemental Movie 6). It appears that mitosis inhibits the chromatin association of Egg, which would prevent it from methylating H3K9.

**Fig. 5.**
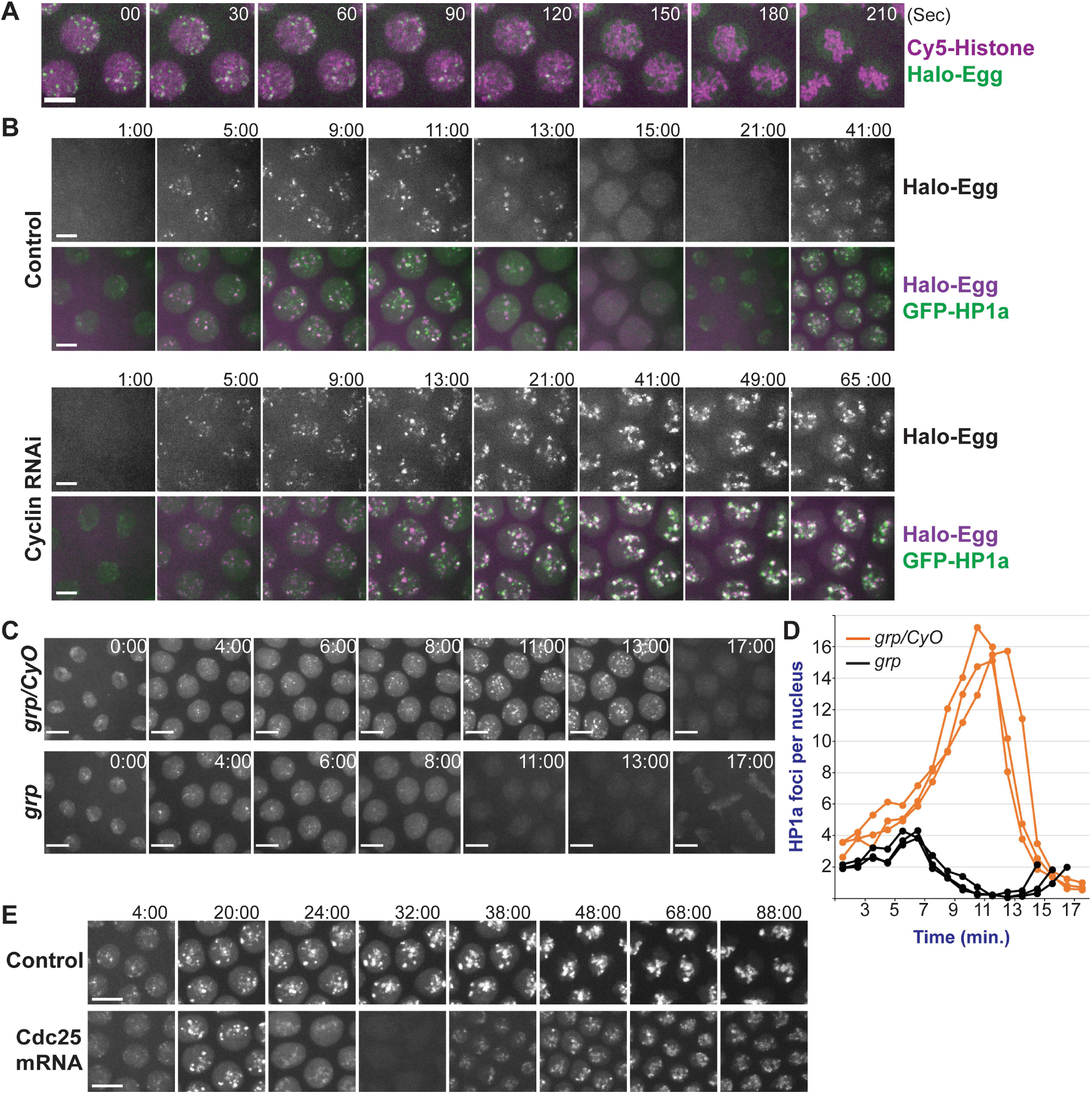
Interphase duration limits the establishment of heterochromatin. **A)** Selected stills from live imaging of Halo-Egg (shown in green) embryos injected with Cy5 labeled Histone proteins (shown in magenta). Egg dissociates from the chromatin upon mitotic chromosome condensation (150 sec.). Scale bar is 3 µm. See Supplemental Movie 6. **B)** Arresting the cell cycle in interphase 13 permits continued accumulation of Egg and HP1a at heterochromatin. Embryos were filmed by confocal microscopy after injection of buffer or dsRNA against cyclins A, B, and B3. Scale bar is 2 µm and the beginning of cycle 13 was set as 0:00. See Supplemental Movie 7. **(C-E)** Shortening interphase duration reduces HP1a accumulation. **C)** Stills from confocal microscopy tracking GFP-HP1a in embryos laid from mothers either heterozygous or homozygous for *grp*. Note that *grp* embryos enter mitosis 13 before completing S phase leading to anaphase bridges (17:00). Scale bar is 4 µm and the start of interphase 13 was set as 0:00. **D)** Plot of the average number of HP1a foci per nucleus over time in control embryos (orange) or *grp* embryos (black). Each line represents data from a different embryo, and foci of GFP-HP1a were counted in approximately 15 nuclei per embryo. **E)** Inducing an early mitosis 14 by expression of twine (cdc25) interrupts heterochromatin formation. Embryos expressing GFP-HP1a were filmed following injection of twine mRNA during cycle 13. Scale bar is 3 µm and the beginning of cycle 14 was set as 0:00.

Our observations over cycles 11-14 (Fig. 3D) when the cell cycle slows naturally suggested that interphase duration limits the recruitment of Eggless to the heterochromatin. To test this idea, we wanted to experimentally arrest the cell cycle in interphase and examine the consequences on Egg localization. Embryos injected with dsRNA against the 3 mitotic cyclins (A, B, and B3) during cycle 11 arrest during interphase 13 (McCleland and O’Farrell 2008). We then followed Halo549-Egg and GFP-HP1a by confocal microscopy during this prolonged interphase. In control embryos, Egg accumulated at nuclear foci over interphase, but by 13 min. into the cycle these foci began disappearing as chromosomes entered mitosis (Fig. 5B). In contrast, arresting nuclei in interphase permitted continued accumulation and chromatin association of Egg for the duration of our records (Fig. 5B, Supplemental Movie 7). Egg foci in arrested nuclei became brighter and larger than in the control and resembled Egg foci observed late in cycle 14 (Fig. 3D). This effect was paralleled by increased recruitment of HP1a to large heterochromatic foci (Supplemental Movie 7). We conclude that mitosis limits Egg accumulation and persistence on chromatin.

Our model would predict that reducing interphase duration would decrease the amount of heterochromatin established by Egg, so we shortened interphase using a grapes (*chk1*) mutant and counted the number of GFP-HP1a foci per nucleus as a readout. Embryos mutant for *grp* have faster cell cycles because the replication checkpoint is required for the progressive prolongation of interphase during cycles 11-13 (Sibon et al. 1997). Mutant *grp* embryos enter mitosis 13 before completing DNA replication and exhibit an 8 min. instead of 14 min. interphase. In heterozygous control embryos the number of HP1a foci increased over the initial 11-12 min. of interphase, reaching a maximum of over 14 foci per nucleus (Fig. 5 C and D), before declining rapidly as nuclei prepare for mitosis. In contrast, in grp embryos HP1a began to recruit to nuclear foci, but the increase in foci number ended abruptly at about 6 min. into interphase (Fig. 5 C and D). Interestingly the rate of increase was also reduced, although the cause of this change is unclear we note that cycles 11 and 12 are also shortened in grp embryos, which could result in a cumulative reduction in the amount of H3K9me2/3 carried over from prior cycles. Thus, shortening interphase reduces the amount of heterochromatin formed.

Injection of *twine* (Cdc25) mRNA during cycle 13 can shorten interphase 14 and lead to an additional mitosis by reintroducing CDK1 activity during a time when it is normally absent (Farrell et al. 2012). We found that the extra mitosis induced in this way interrupted the accumulation of GFP-HP1a (Fig. 5E). Thus, premature mitosis interferes with heterochromatin formation.

## DISCUSSION

Histone modifications play a critical role in the epigenetic programs controlling development. The chromatin in the early *Drosophila* embryo lacks many of these modifications and their restoration begins in an unusual setting where cells divide remarkably quickly and in the absence of substantial transcription. The reappearance of stable chromatin states coincides with the slowing of the cell cycle and the onset of major transcription at the MBT. Here we have focused on the initial events controlling the de novo formation of the heterochromatin. Using our new JabbaTrap method (Fig. 1), we showed that Eggless is the major H3K9 methyltransferase establishing heterochromatin domains in the embryo. Further using real time imaging, we documented the early steps of heterochromatin formation and uncovered a close relationship between the recruitment of Egg (Fig. 3E), the accumulation of H3K9me2/3-HP1a (Fig. 2A) and slowing of the cell cycle. Experiments manipulating interphase duration indicate that this relationship is regulatory (Fig. 5). We propose that the slowing of the cell cycle as the embryo approaches the MBT expands limited windows of opportunity for Egg action and thereby times heterochromatin formation.

### JabbaTrapping – a new tool for inactivating maternal function

The early embryo poses unique challenges to functional studies. The vast maternal supply present for many gene products complicates traditional genetic analysis. Additionally, early embryonic events happen fast so that methods that depend on destruction of RNA and protein must quickly destroy a large pool of target to produce a phenotype. Our JabbaTrap approach can rapidly block maternal activities by mislocalizing target proteins to lipid droplets (Fig. 1). Additionally, by establishing a transgenic system we deployed the JabbaTrap to lipid droplets forming during oogenesis and mislocalized proteins during early embryogenesis. Interestingly, the JabbaTrap also mislocalizes proteins that interact with the target (Fig. 4E) suggesting that it may be a useful approach to inactivate entire nuclear protein complexes. In principle, the JabbaTrap could be applied to any protein with a cognate nanobody, and we observed that the method successfully interferes with many different nuclear functions. However, the function of cytoplasmic proteins might be less susceptible to inhibition by localization to lipid droplets. The combination of nanobody based inactivation approaches and CRISPR-Cas9 gene editing to endogenously epitope tag proteins will allow easy functional studies in the future, and the JabbaTrap fills a special need for such techniques in the dissection of early development.

### A new developmental model for the formation of heterochromatin

The restoration of a more standard cell cycle regulation and timing at the MBT coincides with the formation of mature heterochromatin. However, this is not a single change but a process involving a series of precisely controlled events with the satellite DNA progressively acquiring different defining characteristics of heterochromatin. Direct cytological examination demonstrated that satellite DNA is compacted as early as cycle 8 (Shermoen et al. 2010). Satellites begin exhibiting slight delays in replication time during cycles 10 through 13, and then show major delays during S phase 14. This onset of late replication, and the consequent prolongation of S phase 14 requires the action of the protein Rif1 (Seller and O’Farrell 2018). As the cell cycle slows during the nuclear divisions leading up to the MBT, Egg deposits H3K9me2/3, which recruits HP1a to the heterochromatin. By the end of cycle 14 the satellite sequences are stably marked by H3K9me2/3-HP1a, and they inherit or maintain this modified state throughout development.

Although no obligate coupling is apparent, the embryonic cell cycle program controls the actions of both Rif1 and Egg. Prior to the MBT high levels of cyclin-Cdk1 activity directly inhibit Rif1 by promoting its dissociation from the satellite DNA thus ensuring short S phases (Seller and O’Farrell 2018). The cell cycle itself controls the action of Eggless in two ways (Fig. 6). First, deposition of naïve histones during DNA replication halves the level of histone modification in daughter nuclei. Prior studies have shown that the post S phase recovery of the level of histone modification is slow (Alabert et al. 2015; Xu et al. 2012). Importantly, if a histone-modifying enzyme does not double the amount of its product during a cell cycle then the level of modification will decrease over time. Second, rapid cell cycles limit the opportunity for Egg action. Egg accumulates gradually at chromatin foci during interphase, and accumulated Egg dissociates abruptly when chromosomes condense in preparation for mitosis (Fig. 5A). As interphase extends in the approach to the MBT, both the amount of recruited Egg and its residence time increase, leading to increased deposition of H3K9me2/3. For instance, during the 7-minute interphase 11 Egg does recruit to chromatin, but its accumulation is minimal due to the rush to mitosis (Fig. 3E). Only incremental increases in methyl marks occur prior to the much-elongated interphase 14. Indeed, the large 359 bp repeat only becomes stably associated with H3K9me2/3-HP1a during cycle 14 (Yuan and O’Farrell 2016), and this association requires Egg (Fig. 2). Importantly, experimentally arresting the cell cycle in interphase permitted continued accumulation of Egg and HP1a (Fig. 5B) while shortening interphase reduced the recruitment of HP1a (Fig. 5 C and D). We conclude that the speed of early cell cycle limits the activity of maternally provided Egg and that the schedule of cell cycle lengthening times the appearance of H3K9me2/3-HP1a to give the observed weak accumulations in the later pre-MBT cycles and more fully elaborated heterochromatin at the MBT (Fig 6).

**Fig. 6.**
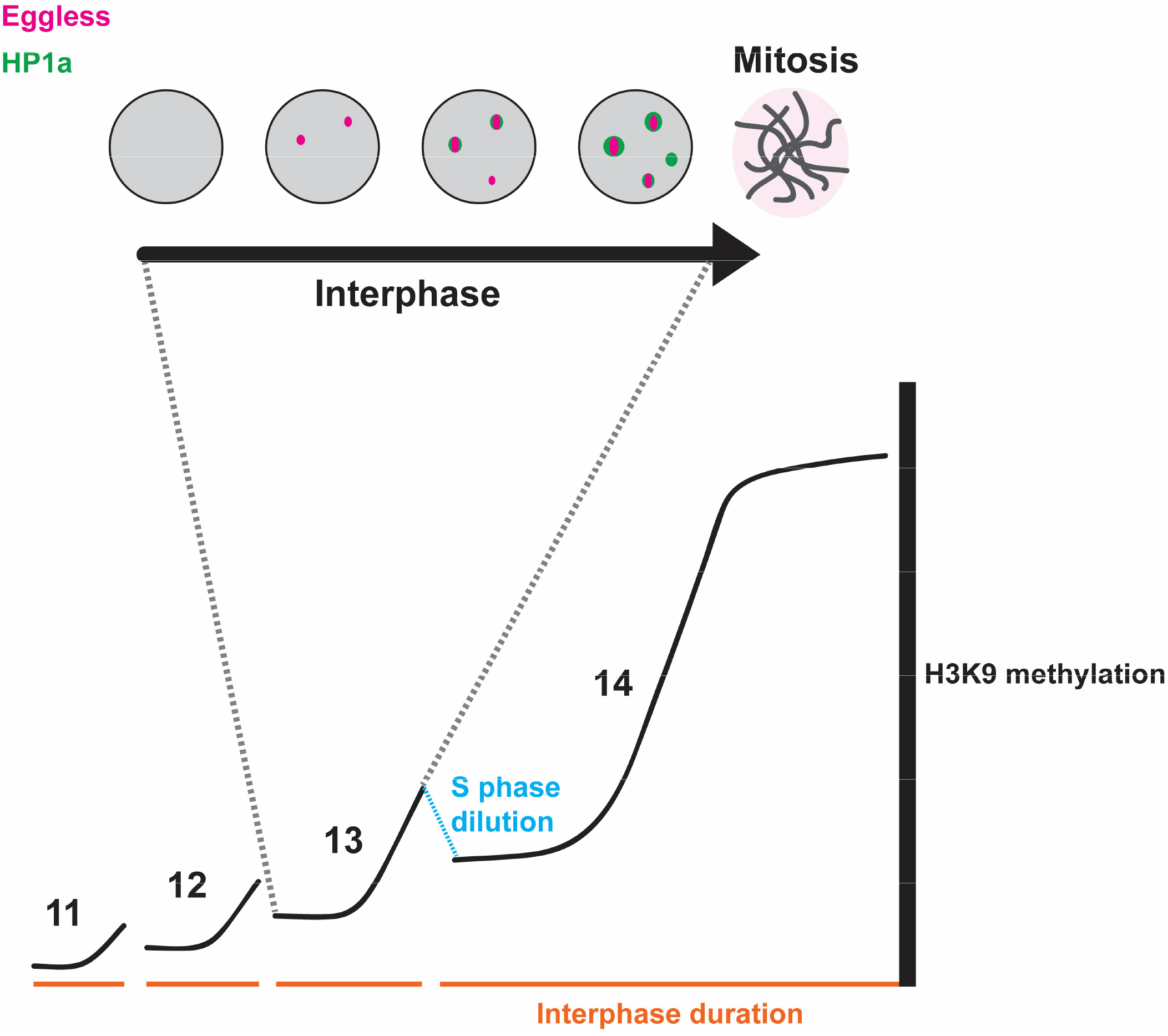
Model. Eggless initiates constitutive heterochromatin formation during early embryonic development. Egg is recruited to chromatin foci after a delay at the start of interphase. Egg foci grow in number and intensity as interphase proceeds. Later in interphase HP1a is recruited to sites occupied by Egg, although some HP1a recruitment is independent of Egg. The domain of HP1a grows and extends beyond the Egg signal. Finally, chromosome condensation in mitosis strips Egg from chromatin. Below a schematic showing that that the progressive accumulation of H3K9-methyl depends on the progressive lengthening of interphase during embryogenesis. Every S phase decreases the level of modification by half due to incorporation of newly synthesized histone proteins. Interphase duration limits the deposition of K9me2/3 by restricting the binding of Egg to its chromatin targets, therefore high levels of modification appear only later in embryogenesis when the cell cycle slows down.

Two recent studies reported that HP1 proteins display liquid like properties, and suggested that the formation of heterochromatin domains depends on the ability of HP1 to phase separate (Larson et al. 2017; Strom et al. 2017). In contrast, our previous observations (Shermoen et al. 2010; Yuan and O’Farrell 2016) suggest that specific interactions trigger the deposition of heterochromatin marks on pre-existing compacted foci of satellite sequences. Here we extend this view by uncovering a role for Egg in the early events of heterochromatin formation. Among methyltransferases, Egg is uniquely required for initiation (Fig. 2). Additionally, using live imaging we documented the first steps in the natural establishment of heterochromatin. Early Egg accumulation localized to small puncta within larger foci of satellite sequences, suggesting a nucleation process. Once recruited to chromatin, Egg deposits H3K9me2/3, and this mark seeds a domain of HP1a, which then grows and spreads perhaps then utilizing HP1’s liquid-like properties (Fig. 6). At least for some of the foci, the small Egg focus does not seem to expand in concert with methylation and HP1a recruitment (Fig. 3A), suggesting additional inputs.

We previously found that timely recruitment of ectopically introduced HP1a to the 359 satellite was blocked by a mutation in its chromoshadow domain that prevents binding to a PxVxL motif (Yuan and O’Farrell, 2106), a seemingly unexpected dependency. This same study found that HP1a bearing a mutation blocking K9me2/3 recognition (V26M mutant) still associated with this satellite, a finding that appears at odds with the present finding that recruitment of HP1 to 359 requires the methyltransferase Egg. The meaning of these observations is not clear. Perhaps dimerization or oligomerization allowed the ectopically introduced V26M mutant to localize with endogenous wild-type HP1a that was recruited to foci of H3K9me2/3. Alternatively, Egg might have methylation independent interactions that can recruit other proteins including HP1a in a heterochromatin complex. The findings suggest that complex interactions guide maturation of heterochromatin foci, and many details remain to be uncovered. Importantly, our present study identifies Egg as a key factor in the earliest steps of this process.

Given that we observed recruitment of Egg to specific chromatin foci as early as cycle 10 (Fig. 3D) the inputs specifying the future heterochromatic sequences must be present very early. However, we can currently only speculate about how Egg is targeted. One attractive hypothesis is that specificity comes from maternally deposited small RNAs that would recognize complementary nascent transcripts, much like the heterochromatin machinery of S. pombe (Allshire and Madhani 2018). In support of this notion the recruitment of HP1a to the 359 repeat requires the maternal presence of this satellite indicating that this process is maternally guided (Yuan and O’Farrell 2016). Additionally, Egg participates in the Piwi pathway and is directed to transposons in the genome by complementary piRNAs, although the details of this mechanism are currently unclear (Sienski et al. 2015). A second possibility is that Egg binds to specific DNA binding proteins that act to specify its localization. A family of DNA binding proteins known as KRAB-ZFP repressors target SetDB1 in mammals (Schultz et al. 2002). While the KRAB family appears to be a tetrapod specific innovation, proteins recognizing specific satellite repeats have been identified in *Drosophila*, and it remains a possibility that they recruit Egg to chromatin (Aulner et al. 2002).

### Developmental specificity of methyltransferase action

Development can employ the same protein for different purposes in different tissues and stages, and Egg is prime example of this. Our results demonstrate that Egg plays a major role in the initial deposition of heterochromatin in the embryo, but prior work had shown that loss of egg does not affect the association of H3K9me2/3-HP1a at the chromocenter in larval polytene chromosomes (Seum et al. 2007a; Tzeng et al. 2007). So Egg is critical for establishment, but not for the maintenance of heterochromatin. Interestingly, the reverse is true for the methyltransferase Su(var)3-9. It makes only minimal contribution during initiation (Fig. 2B and C), but then plays a dominant role later in development. We suggest that one key difference between these two enzymes underlies this developmental specialization. Su(var)3-9, but not Egg, contains a chromodomain allowing it to bind its own product making it ideal for maintenance (Schotta et al. 2002). This capacity allows for positive feedback where prior methylation facilitates the modification of adjacent nucleosomes. In fission yeast the chromodomain of the K9 methyltransferase Clr4 is critical for the spread and maintenance of domains of H3K9me2/3 (Hall et al. 2002; Zhang et al. 2008; Al-Sady et al. 2013). Notably, a recent study in C. elegans embryos uncovered that the SETDB1 methyltransferase (MET-2) is critical in this species for the formation of heterochromatin, indicating that a special function for SETDB1 during initiation is conserved (Mutlu et al. 2018).

### Do rapid embryonic cell cycles wipe the slate clean during development?

Although we have focused on the constitutive heterochromatin, our study could provide broader insight into the regulation of histone modifying enzymes during development. In fact many other histone modifications only appear at the MBT (Chen et al. 2013; Li et al. 2014), and direct examination of the enzymes installing these marks during embryogenesis would be informative. We have emphasized the limitations that cell cycle speed imposes on the restoration of the histone modifications, but we don’t want to obscure what we think may be an important benefit of these rapid divisions. Embryos inherit their genomes from their parents, and these parental genomes come with parental chromatin modifications that are installed during gametogenesis (Loppin et al. 2005). The vast majority of animals develop externally from large eggs that begin with rapid cell divisions without growth or transcriptional input. We suggest these fast cell cycles provided an ancestral mechanism to reset the embryonic chromatin without the need for specific enzymes catalyzing the removal of histone modifications. Multiple rounds of short S phases would dilute parental modifications, and brief interphases would limit the action of maternally supplied chromatin modifying enzymes, which are present long before they are needed. Developmental programs could then re-introduce these so-called epigenetics marks as the embryo reaches the MBT, thereby coordinating the onset of zygotic transcription with the appearance of stable patterns of histone modification. Notably, the situation in mammalian embryos is different with repressive modifications appearing early during the cleavage stages (Fadloun et al. 2013). However, the evolution of placental development restructured mammalian embryogenesis and the post-fertilization divisions are much slower and transcription begins immediately. By controlling the speed of the embryonic cell cycle, development coordinates the deposition of epigenetic modifications with the onset of zygotic transcription.

## METHODS

### Fly stocks

Stocks of *Drosophila melanogaster* were cultured on standard cornmeal-yeast media. The following previously described stocks were used in this study: *w^1118^* Canton-S (wild type), *His2Av-mRFP* (Bloomington stock number 23650), *EGFP-HP1a* (30561), *maternal tubulin-Gal4* (7063), *sfGFP-zelda* (M. Harrison), *ORC2-GFP; orc2^1/y4^* (M. Gossen), *G9a^RG5^* (P. Spierer), *grp^06034^, Su(var)3-9^06^, Su(var)3-9^17^*, and *RIF1-EGFP*.

Embryo injections for the transgenic lines described in this study were done by Rainbow Transgenic Flies in Camarillo, CA. PhiC31 mediated integration generated the JabbaTrap transgene *UASp-VhhGFP4-jabba-VhhGFP4^attP-9A^*. The following stocks were created using CRISPR-Cas9 gene editing: *sfGFP-Su(z)12, sfGFP-G9a, EGFP-egg*, and *Halo-egg*. See Supplemental Fig. 1 for details of endogenous tagging. Following generation, modified flies were crossed to our lab’s wild type stock (*w^1118^ Canton-S*) for at least 5 generations. Plasmid DNA and sequence files are available upon request.

### Protein purification

Purified HP1a and TALE-light GFP/mCherry fusion proteins were described in previous studies (Shermoen et al. 2010; Yuan and O’Farrell 2016). Halo tagged 359 TALE-light protein was produced in BL21(DE3) *E. coli* cells. Protein expression was induced by treating cultures with 0.5 mM IPTG overnight at room temperature. Bacteria were lysed using B-PER supplemented with lysozyme, DNase I, and PMSF. TALE-light proteins were purified from cleared lysate using Ni-NTA agarose in gravity flow columns. After 2 washes, 10 µM of the halo ligand JF646 in wash buffer was added to label the protein on column. Labeled proteins were eluted, dialyzed into 40 mM HEPES (pH 7.4) with 150 mM KCl, and then concentrated by spin column. The concentrated eluate was supplemented with glycerol to 10% and then snap frozen in liquid N_2_.

### In vivo Halotag labeling of Drosophila embryos

We adapted the embryo permeabilization method described in (Rand et al. 2010) to deliver the Halotag ligand into the embryo, and permeabilization and labeling were conducted in a single step. Embryos were collected in mesh baskets, dechorinated with 50% bleach, and then rinsed thoroughly with Embryo wash buffer and then water. Using a brush, embryos were transferred to microfuge tubes with a 1:10 solution of Citrasolv containing 1 µM JF549. The tubes were incubated on a nutator in the dark for 8 to 9 minutes. Embryos were then poured into mesh baskets and cleaned thoroughly with wash buffer and water to remove excess detergent. Embryos were then prepared for imaging following our normal protocol. Permeabilization was not always successful, but embryos containing the dye could easily be identified microscopically.

### Immunoprecipitation and Western blotting

Immunoprecipitation and western blotting were performed as described in (Seller and O’Farrell, 2018). The following primary antibodies were used at a dilution of 1:1000: rabbit anti-GFP (ab290, Abcam, Cambridge, United Kingdom) and rabbit anti-Wde (gift from A. Wodarz). A Goat anti-Rabbit HRP conjugated secondary antibody (Biorad) was used at a dilution of 1:10,000.

### Embryo immunostaining

Embryos were collected on grape agar plates, washed into mesh baskets and dechorinated in 50% bleach for 2 minutes. Embryos were then devitellenized and fixed in a 1:1 mixture of methanol-heptane, before storing in methanol at −20°C. Embryos were gradually rehydrated in a series of increasing PTx:Methanol mixtures (1:3, 1:1, 3:1) before washing for 5 minutes in PTx (PBS with 0.1% Triton). Embryos were then blocked in PTx supplemented with 5% Normal Donkey Serum and 0.2% Azide for 1 hour at room temperature. Blocked embryos were then incubated with the primary antibody overnight at 4°C. The following primary antibodies were used: rabbit anti-Wde at 1:500 (A. Wodarz), rabbit anti-H3K9me3 at 1:500, mouse anti-Ubx at 1:20 (DSHB FP3.38) and mouse anti-HP1a at 1:100 (DSHB CA19). Embryos were then washed with PTx for three times 15 minutes each, and then incubated with the appropriate fluorescently labeled secondary antibody (Molecular Probes) at 1:500 for 1 hour in the dark at room temperature. Embryos were then washed again with PTx for three times 15 minutes each. DAPI was added to the second wash. Finally, stained embryos were mounted on glass slides in Fluoromount.

### Embryo microinjection and microscopy

Embryo microinjections were performed as described in (Farrell et al. 2012). JabbaTrap and Cdc25 (twine) mRNAs were synthesized using the CellScript T7 mRNA production system and injected at a concentration of 600 ng/µl. dsRNA against the cyclins A, B, and B3 was prepared as described in (McCleland and O’Farrell 2008). Imaging was performed using a spinning disk confocal microscope as described in (Seller and O’Farrell 2018). Image processing and analysis were done using Volocity (Perkin Elmer) and ImageJ. Unless otherwise noted, all images are presented as z-stack projections.

## Acknowledgements

We thank past and present members of the O’Farrell laboratory, especially Kai Yuan, Antony Shermoen, and Mark McCleland for guidance and reagents. We also thank Anna Holman for early work on this project. Hiroshi Kimura generously provided labeled Fab reagents for H3K9me2. Luke Lavis generously provided halotag ligands labeled with Janelia Fluorescent dyes. The Bloomington Stock Center and the *Drosophila* community provided important reagents and fly stocks throughout this work.

## Author Contributions

**Charles Seller:** Conceptualization, Investigation, Writing.

**Chun-Yi Cho:** Investigation, Editing.

**Patrick O’Farrell:** Conceptualization, Writing, Supervision.

